# Vaginal and uterine microbiomes in beef cattle at artificial insemination and associations with pregnancy outcomes

**DOI:** 10.64898/2026.03.31.715609

**Authors:** Justine Kilama, Devin B. Holman, Joel S. Caton, Kevin K. Sedivec, Carl R. Dahlen, Samat Amat

## Abstract

The female reproductive tract harbors complex microbial communities that may influence reproductive success. In previous work using 16S rRNA gene sequencing, we identified bacterial taxa in the vagina and uterus of beef cattle associated with pregnancy outcomes, but taxonomic resolution and functional inference was limited. Here we used shotgun metagenomic sequencing to characterize the taxonomic composition, functional potential, and antimicrobial resistome of vaginal and uterine microbiomes at the time of artificial insemination (AI) in cows that subsequently became pregnant or remained open. Vaginal (pregnant n = 54; open n = 7) and uterine (pregnant, n = 41; open, n = 9) samples were collected prior to AI. Microbial community structure did not differ between pregnancy outcome groups in either anatomical site (PERMANOVA; *P* > 0.05). However, cows that remained open showed significantly greater species-level richness and diversity in the vaginal microbiome (*P* < 0.05). No diversity differences were observed in the uterine microbiome. In contrast, significant differences were detected between anatomical sites, with distinct dominant taxa and functional profiles. Vaginal microbiomes were enriched in pathways related to genetic information processing, whereas uterine microbiomes exhibited greater representation of metabolic pathways. A total of 105 ARGs spanning 11 antimicrobial classes were identified, with tetracycline resistance genes [*tet*(Q), *tet*(W), and *tet*(M)] predominating, and bla_TEM-116_ more abundant in the uterine microbiome. Overall, while vaginal and uterine microbiomes were compositionally and functionally distinct, no robust pregnancy-associated taxonomic or functional signatures were detected, likely reflecting limited statistical power and challenges inherent to low-biomass metagenomic datasets.

**IMPORTANCE:** Understanding the role of the reproductive tract microbiome in fertility could improve reproductive efficiency in cattle. We used shotgun metagenomic sequencing to characterize the taxonomic composition, functional potential, and antimicrobial resistome of vaginal and uterine microbiomes at the time of artificial insemination in cows that subsequently became pregnant or remained open. Using paired samples from the same animals, we directly compared microbial communities between the upper and lower reproductive tract to identify shared and site-specific features. Although no distinct microbial signatures associated with pregnancy outcomes were detected, this may reflect limited statistical power and low microbial biomass inherent to these samples. Despite these challenges, our study provides high-resolution insights into the composition, functional potential, and resistome of bovine reproductive microbiomes and highlights important technical considerations for studying low-biomass microbial ecosystems.

## INTRODUCTION

Reproductive inefficiency remains a critical challenge hindering herd productivity and reducing profitability in beef and dairy cattle systems in the United States (Reese et al., 2020). A recent meta-analysis estimated that roughly 48% of beef cows experience pregnancy loss within the first month following a single AI, with an additional 6% occurring later in gestation (Reese et al., 2020). These reproductive setbacks extend calving intervals, increase rebreeding costs, and reduce the number of calves weaned, representing a major economic burden for cattle producers (Lamb et al., 2010; Reese et al., 2020). Despite advancements in genetic selection, nutrition, and reproductive management, improvements in timed AI success rates have been modest over recent decades, suggesting that important biological drivers of conception and early gestation remain poorly understood (Berry et al., 2014; Diskin et al., 2016).

The mammalian reproductive tract, particularly the vagina and uterus, harbors complex and dynamic microbial communities that are important in maintaining reproductive health and can influence fertility outcomes (Baud et al., 2018; Ault et al., 2019; Baud et al., 2019; Koedooder et al., 2019a; Koedooder et al., 2019b; Baud et al., 2023b; Webb et al., 2023). In humans, vaginal microbiome composition has been linked to pregnancy likelihood in both natural conception and assisted reproductive procedures, such as *in vitro* fertilization and embryo transfer (Koedooder et al., 2019b; Odendaal et al., 2024). In cattle, high-throughput sequencing has identified distinct microbial communities in the uterus, challenging the long-held assumption of uterine sterility (Moore et al., 2017; Ault et al., 2019). Recent studies further indicate that reproductive success in cattle may depend on a complex interplay of factors, including the urogenital tract microbiota (Ong et al., 2021; Luecke et al., 2022; Webb et al., 2023). Dysbiosis in these microbial communities has been associated with diseases such as metritis and endometritis (Silva et al., 2024) (Jeon et al., 2015; Galvão et al., 2019), which are often characterized by reduced microbial diversity and an overrepresentation of potential pathogens, including *Bacteroides, Fusobacterium, Porphyromonas,* and *Trueperella* spp. (Galvão et al., 2019; Ong et al., 2021; Çömlekcioğlu et al., 2024). These observations raise the possibility that even subtle changes in the reproductive tract microbiota may influence fertilization, embryo implantation, and early gestational development in cattle (Adnane and Chapwanya, 2022).

In our previous study using 16S rRNA gene sequencing, we identified microbial taxa in the vaginal and uterine tracts at the time of AI that were positively associated with pregnancy success (Webb et al., 2023). A total of 11 vaginal taxa belonging to the phylum Bacillota, including members of the *Lachnospiraceae Oscillospiraceae* families, were significantly enriched in open heifers as compared to those that become pregnant following AI (Webb et al., 2023). Similarly, 28 uterine bacterial taxa were differentially abundant between open and pregnant group cows, with 11 taxa including the *Faecalibacterium* and *Fusobacterium* spp. being enriched in pregnant cows (Webb et al., 2023). However, 16S rRNA gene sequencing is inherently limited in taxonomic resolution, often restricting identification to the genus level, and does not directly assess functional capacity (Langille et al., 2013; Moore et al., 2017; Walsh et al., 2018). Therefore, in this study, we used shotgun metagenomic sequencing to identify taxonomic and functional signatures of vaginal and uterine microbiomes at the time of AI that are associated with pregnancy outcome. As a secondary objective, we profiled antimicrobial resistance genes (ARGs) in the vaginal and uterine microbiomes, considering the bovine reproductive tract as a potential reservoir and transmission route, with implications for the transfer of ARGs to bulls and offspring.

## MATERIALS AND METHODS

All animal procedures were conducted in accordance with the guidelines of the North Dakota State University Institutional Animal Care and Use Committee (IACUC; protocol #A21061).

### Animal husbandry and experimental design

The experimental design, animal management, estrus synchronization, and swab collection procedures followed protocols described in our previous work (Webb et al., 2023). In this follow-up study, we investigated the reproductive tract microbiome of Angus-crossbred beef cows (N = 100) with a mean age of 5.49 ± 2.77 years (Figure 1). Vaginal and uterine swabs were simultaneously collected from each cow immediately prior to AI, as previously outlined (Webb et al., 2023). Briefly, cows underwent estrus synchronization using the 7-day Co-Synch + CIDR (controlled internal drug release) protocol, which involved the insertion of a CIDR progesterone intravaginal device (EaziBreed CIDR, Zoetis, Parsippany, NJ, USA). On day 0, cows received an injection of gonadotropin-releasing hormone (GnRH) followed immediately by CIDR insertion, which remained in place for 7 days. Upon CIDR removal, cows were administered prostaglandin to induce estrus. Fixed-time AI was performed 60–66 h later, along with a second injection of GnRH.

**Figure 1:**
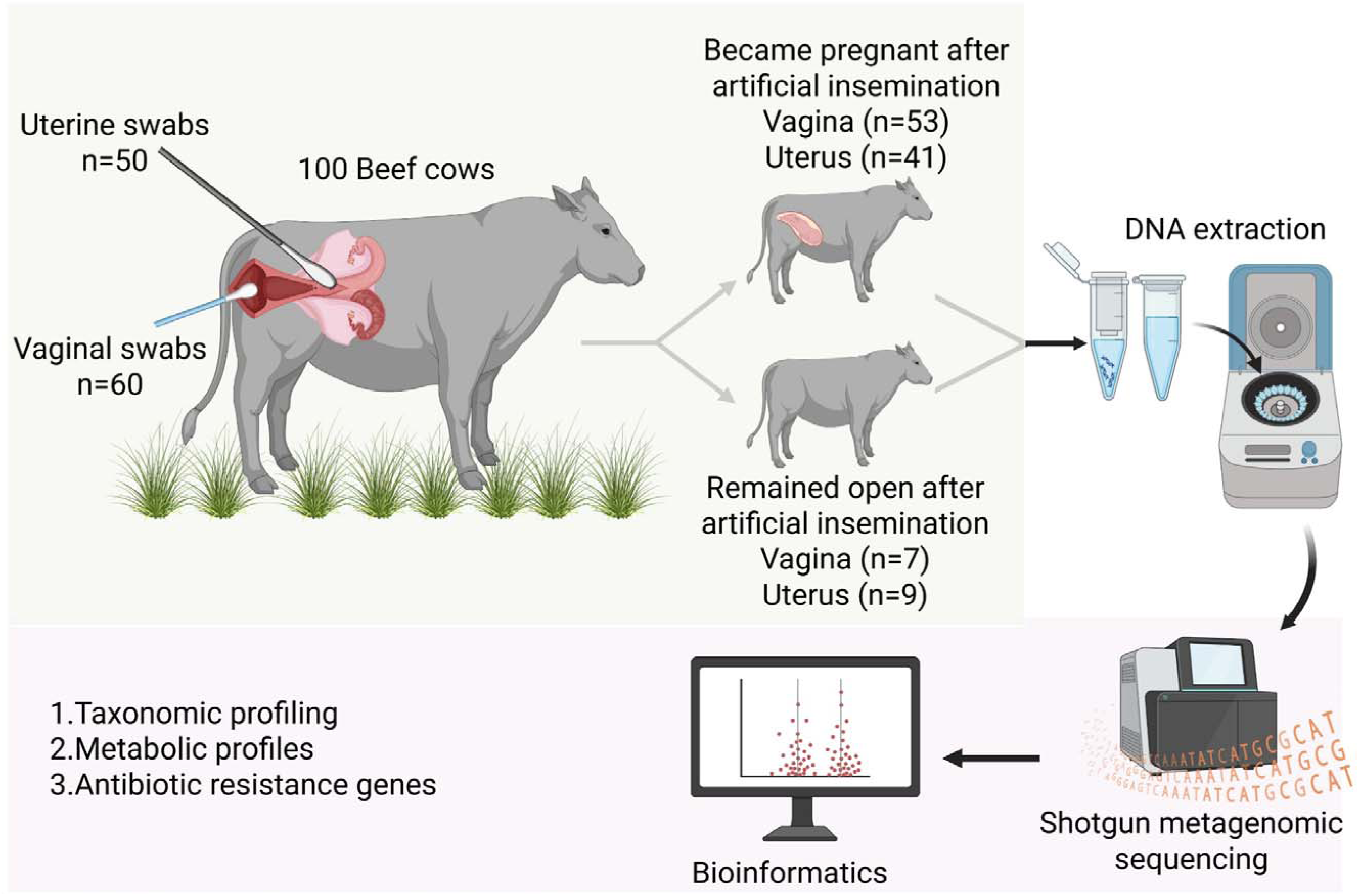
Schematic representation of the experimental design, sampling strategy, and sample processing workflow. Vaginal and uterine swabs were collected from 100 Angus beef cows 2 days prior to artificial insemination. A subset of swabs from each anatomical site were selected for downstream processing: vaginal (n = 60) and uterine (n = 50). Samples were categorized based on pregnancy outcome determined at 35 days post-AI: pregnant (vaginal, n = 7; uterine, n = 9) and open (vaginal, n = 53; uterine, n = 41). DNA was extracted from the swabs and subjected to shotgun metagenomic sequencing. Downstream bioinformatics analyses included removal of the *Bos taurus* reference genome, taxonomic classification of metagenomic reads, metabolic pathway profiling, and identification of antibiotic resistance genes (ARGs).

All cows were approximately 84 days postpartum, were suckled, and had been pasture-grazed for 45 days prior to sampling, with no postpartum disease observed. During summer 2023, the pasture at the Central Grasslands Research Extension Center (CGREC) was classified as mixed-grass prairie dominated by western wheatgrass (*Pascopyrum smithii*), green needlegrass (*Nassella viridula*), and blue grama (*Bouteloua gracilis*)(McCarthy et al., 2023). Pregnancy status was determined at 35 days post-AI using an ExaGo ultrasound system (IMV Technologies, France) equipped with an IMV 13 cm, 3.5 MHz linear transducer, and animals were classified as either pregnant or open.

### Vaginal and uterine swab collection

Vaginal samples were collected from cows (n = 100) at the time of AI, following procedures described previously (Webb et al., 2023). The vulva was disinfected with 70% ethanol, and a sterile 15 cm cotton swab (Puritan, Guilford, ME, USA) was gently inserted into the vaginal cavity. Once positioned in the center of the vaginal canal, the swab was rotated four times to ensure adequate sampling and then slowly withdrawn to minimize contamination. Swabs were immediately placed into sterile Whirl-Pak bags (Uline, Pleasant Prairie, WI, USA), kept on ice, and transported to the laboratory for processing.

Uterine samples were collected from these 100 cows immediately after vaginal swabbing using a 71-cm double-guarded swab (Reproduction Provisions L.L.C., Walworth, WI, USA) according to the method we described previously (Webb et al., 2023). Briefly, guided by rectal palpation, the swab was passed through the vagina and cervix into the uterine body, where the tip was extended and rotated three times with gentle pressure. The swab was then retracted, removed, and placed into a 2-mL tube on ice for transport to the laboratory. Vaginal and uterine swabs were collected concurrently from each cow within a 4-hour window by the same personnel. In the laboratory, swabs were transferred into 2-mL screw-cap tubes containing 1 mL of brain heart infusion (BHI) broth (Hardy Diagnostics, Santa Maria, CA, USA) with 20% glycerol (Fisher Scientific, Fair Lawn, NJ, USA) and stored at −80°C until further processing.

### Sample selection for metagenomic DNA extraction

Most cows in this cohort became pregnant, leaving only 13 non-pregnant animals available for inclusion as open cows. Preference was given to those animals who had paired vaginal–uterine samples, which limited the number of open cow samples available from both sites.

### Genomic DNA extraction from vaginal and uterine swab samples

Genomic DNA was extracted from vaginal swabs using the DNeasy Blood and Tissue Kit (Qiagen Inc., Hilden, Germany), following the protocol outlined in our previous study (Webb et al., 2023). Briefly, thawed samples were centrifuged to pellet the microbial cells, and swab tips were cut and added to the corresponding pellets. Samples were then subjected to enzymatic lysis, proteinase K digestion, and two rounds of bead beating. DNA was purified using silica columns, eluted in 50 µL of pre-warmed buffer, and quantified with a NanoDrop ND-1000 spectrophotometer and PicoGreen assay. A sterile swab processed in parallel served as a negative extraction control. DNA was stored at -20°C until being sent for metagenomic library preparation.

Genomic DNA was extracted from uterine swabs (41 pregnant and 9 open) using a modified cetyltrimethylammonium bromide (CTAB) and phenol:chloroform protocol, as described previously (Fujimura et al., 2016; Rackaityte et al., 2020; Amat et al., 2021). This extraction method was selected because preliminary trials yielded higher-quality DNA compared to the commercial kits used in our earlier 16S rRNA gene sequencing study. Uterine swabs stored in BHI broth with 20% glycerol were vortexed, and 500 µL of suspension was transferred into 2-mL screw-cap tubes. After centrifugation at 16,000 × *g* for 30 min at 4°C, pellets were resuspended in 500 µL of pre-warmed CTAB buffer, transferred to Lysing Matrix E tubes, and subjected to thermal lysis (65°C, 15 min) followed by bead beating (5.5 m/s, 30 s). DNA was extracted with phenol:chloroform:isoamyl alcohol (25:24:1), cleaned with chloroform in phase-lock gel tubes, and precipitated with linear acrylamide and PEG-NaCl. DNA pellets were washed with 70% ethanol, eluted in 30 µL of 10 mM Tris-Cl (pH 8.0), quantified with a NanoDrop ND-1000, and stored at −20°C. A sterile nuclease-free water control was included as a negative extraction control throughout the DNA extraction process.

### Library preparation and shotgun metagenomic sequencing

Genomic DNA samples were first subjected to quality control to assess concentration and integrity prior to library construction. From amongst the DNA samples processed, 60 vaginal and 51 uterine samples were selected for library preparation and sequencing. Shotgun metagenomic sequencing libraries were prepared and sequenced by Novogene (Sacramento, CA, USA). In brief, DNA was randomly fragmented and processed through end-repair, A-tailing, and ligation of Illumina sequencing adapters. Adapter-ligated fragments were size-selected and PCR-amplified to generate sequencing libraries. Library concentration was quantified using a Qubit fluorometer and qPCR, and fragment size distribution was assessed using a fragment analyzer. Libraries were pooled and sequenced on an Illumina NovaSeq X Plus platform (Illumina Inc., San Diego, CA, USA) using paired-end 150 bp reads (PE150). Raw sequencing reads were quality filtered to remove reads containing adapter contamination, reads with more than 10% ambiguous bases (N), and reads containing more than 50% low-quality bases (Q ≤ 5).

### Metagenomic sequencing data analysis

Sequencing adapters were trimmed from the raw metagenomic reads, and low-quality sequences were filtered using fastp v.0.24.1 (Chen et al., 2018), with a 4-bp sliding window and a minimum quality score of 15. Reads shorter than 100 bp were discarded. The remaining reads were aligned to the *Bos taurus* reference genome (ARS-UCD2.0) and the *Escherichia phage* phiX174 (NC_001422) using Bowtie2 v.2.5.4 (Langmead and Salzberg, 2012). SAMtools v.1.22.1 (Danecek et al., 2021) and BEDtools v.2.31.1 (Quinlan and Hall, 2010) were used to extract reads that did not map to these genomes. A second round of mapping was performed, incorporating the reference genome for *Bos indicus* (NIAB-ARS_B.indTharparkar_mat_pri_1.0) and the human genome (GRCh38.p14).

Taxonomic classification was performed using Kraken2 v.2.1.6 (Wood et al., 2019) and Bracken v.3.1 (Lu et al., 2017) with a minimum read count threshold of 10, using the Genome Taxonomy Database (GTDB) release 226 (Parks et al., 2022). To minimize host misclassification, the GTDB was supplemented with the *B. taurus, B. indicus*, and human genomes. Reads that Bracken redistributed to host genomes were removed, and the relative abundance of archaeal and bacterial taxa was calculated as the number of reads assigned to each taxon divided by the total number of reads assigned to archaea and bacteria.

### Antimicrobial resistome profiling

The metagenomic reads were also screened for antimicrobial resistance genes (ARGs) using the Resistance Gene Identifier (RGI) v.6.0.5 and the Comprehensive Antibiotic Resistance Database (CARD) v.4.0.1 (Alcock et al., 2023) with KMA (Clausen et al., 2018). The functional gene content was assessed by aligning the metagenomic reads to the Kyoto Encyclopedia of Genes and Genomes (KEGG) release 114.0 (Kanehisa et al., 2023) prokaryotic protein database with DIAMOND v.2.1.8.162 (Buchfink et al., 2021) and assigning genes to KEGG orthology (KO) groups.

### Bioinformatics and statistical analysis

For both the uterine and vaginal microbiomes, permutational multivariate analysis of variance (PERMANOVA) based on Bray-Curtis dissimilarities was conducted using relative abundance data for archaeal and bacterial genera and species, as well as copies per million reads for KOs, to assess overall compositional differences between cattle that became pregnant and those that remained open. Shannon diversity and inverse Simpson diversity indices were also calculated using the relative abundance values at the genus and species level. All analyses were conducted in R v.4.5.0 using the vegan package (v.2.7-1). MaAsLin3 v.1.0.0 (Nickols et al., 2026) was used to identify differentially abundant microbial species and genera as well as KOs and KEGG pathways in the uterine and vaginal microbiomes of cattle that either became pregnant or remained open. Only those microbial species and KEGG pathways found in at least 25% of the samples analyzed, and microbial species with a relative abundance of ≥ 0.01%, were included for the differential abundance analyses (53 vaginal and 43 uterine samples).

For the alpha diversity measures, normality was assessed using the Shapiro-Wilk test, and Levene’s test was used to evaluate the homogeneity of variances. Due to violations of normality assumptions, non-parametric Mann–Whitney U tests were conducted in SAS version 9.4 (SAS Institute Inc., Cary, NC, USA) to compare alpha diversity indices between pregnant and open cows within the vaginal and uterine microbiomes. Statistical significance was defined as *P* < 0.05. Data visualizations were generated using GraphPad Prism and the ‘vegan’ package in R.

## RESULTS

### Metagenomic sequencing summary and host reads removal

Prior to processing, there were 47,751,145 ± 882,211 (mean ± SEM; n = 60) reads for vaginal samples and 85,678,320 ± 2,145,365 reads (mean ± SEM; n = 51) for the uterine samples. The higher read count for uterine samples reflects that these libraries were sequenced twice and subsequently combined. Due to the low ratio of microbial biomass to host DNA, two rounds of host read removal were performed. Following processing, the number of reads was reduced to 218,934 ± 140,307 reads per vaginal sample and 205,798 ± 16,558 reads per uterine sample. One vaginal sample (B46; 8,633,418 reads) contained more than 10 times as many post-processing reads as any other sample (Table S1). To avoid bias, this sample was excluded from downstream taxonomic, functional, and ARG analyses.

Initially, the taxonomic composition of these samples was profiled using the unmodified GTDB, which included only archaeal and bacterial genomes. When a subset of reads initially classified as bacterial by Kraken2 was randomly selected and analyzed with BLASTn, many appeared to instead be from *B*. *taurus* or *B*. *indicus* (data not shown). This suggests that two rounds of host DNA removal may not have been sufficient and that residual host-derived reads may have been misclassified as bacterial. Such misclassification has also been reported previously, as unaccounted host DNA can be incorrectly assigned as microbial sequences by Kraken2, particularly in low-biomass samples (Gihawi et al., 2023). As recommended by Gihawi et al. (2023), we supplemented the GTDB with *B*. *indicus* and *B*. *taurus* genomes, as well as the human genome, to account for potential human contamination during sampling, DNA extraction, or library preparation. As a result, a number of reads previously classified as bacterial were reassigned to *B*. *indicus*, *B*. *taurus*, *Homo sapiens*, or left unclassified.

### Vaginal microbiome

The overall vaginal microbial community structure at the genus level did not differ significantly between cows that became pregnant and those that remained open (Fig. S1A; *R^2^* < 0.001, *P* = 0.197). Similarly, no significant differences (*P* > 0.05) were observed in genus-level alpha diversity metrics (Fig. S1B-D). However, at the species level, cows that remained open exhibited significantly greater species richness (Fig. 2A; *P* = 0.0427) and higher microbial diversity, as measured by the inverse Simpson index (Fig. 3B; *P* = 0.0299) and Shannon diversity index (Fig. 2C; P = 0.031). The vaginal microbiome at the genus level was dominated by *Streptococcus* (26.17%) and *Ureaplasma* (20.74%) (Fig. 3A). At the species level, the most relatively abundant taxa were *Streptococcus pluranimalium* (33.89%) and *Ureaplasma diversum* (26.84%) (Fig. 3B). No genera or species were differentially abundant in the vaginal microbiome between open and pregnant cows (*P* > 0.05). Two samples (B56, B58) had no sequences confidently assigned (≥ 10 reads) to a unique species and therefore were excluded from the relative abundance calculations.

**Figure 2.**
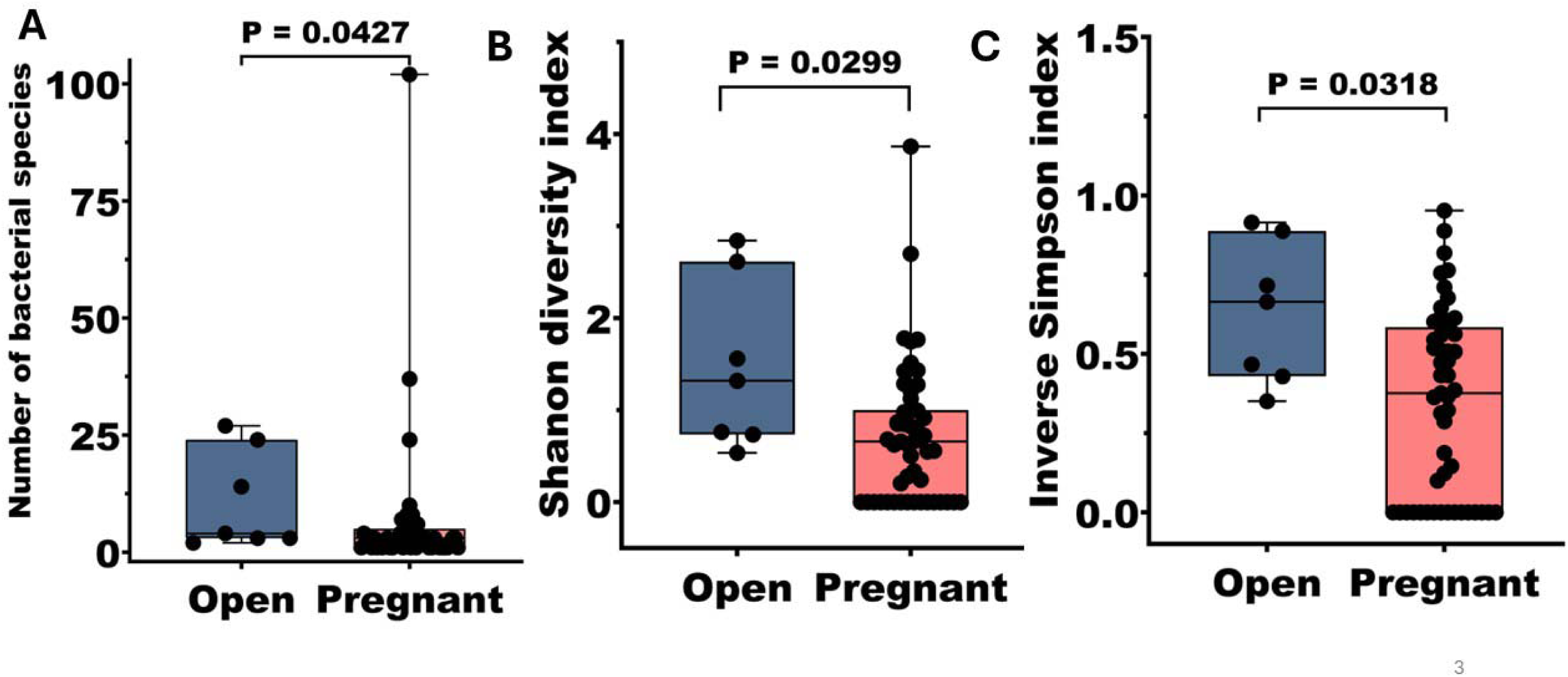
Alpha diversity of the vaginal microbiome at the species level in cows that remained open (n = 7) or became pregnant (n = 53) following artificial insemination. Box-and-whisker plots showing alpha diversity metrics, including **(A)** observed species richness**, (B)** Shannon diversity index, and **(C)** inverse Simpson diversity index. Boxes represent the interquartile range (IQR), the horizontal line indicates the median, and whiskers extend to 1.5 × IQR. Individual data points are shown as jittered dots. Statistical differences in species richness and alpha diversity of the vaginal microbiome between cows that remained open (n = 53) or became pregnant (n = 7) following artificial insemination were evaluated using the Mann-Whitney U test (two-tailed), with significance declared at P < 0.05.

**Figure 3.**
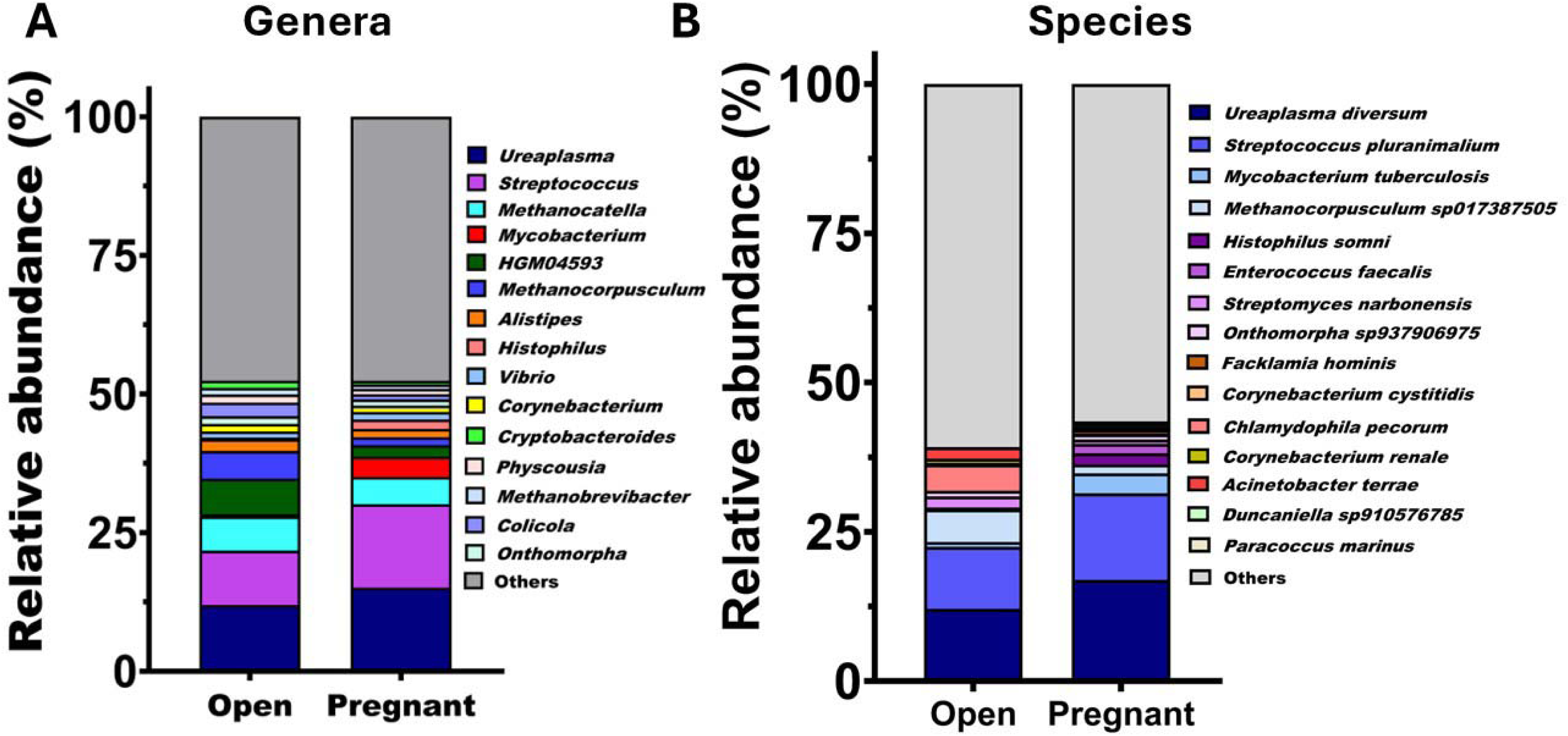
Stacked bar charts showing taxonomic composition of the vaginal microbiome in cows that remained open (n = 7) or became pregnant (n = 53) following artificial insemination. **(A)** Relative abundance (%) of the 15 most abundant bacterial genera; **(B)** Relative abundance (%) of the 15 most abundant bacterial species.

Functional profiling based on KEGG pathways showed that nucleotide metabolism was the most abundant functional category in the vaginal microbiome, followed by membrane transport, replication and repair, translation, and energy metabolism (Fig. 4). Other notable categories included carbohydrate metabolism, and amino acid metabolism. The most abundant individual pathways were ABC transporters, aminoacyl-tRNA biosynthesis, purine metabolism, glycolysis/gluconeogenesis, and ribosome (Fig. 4). Although cows that remained open (n = 6) showed a trend toward higher mean copies per million reads (CPM) across functional categories than pregnant cows (n = 47), no pathways differed significantly between groups (Fig. S3A).

**Figure 4.**
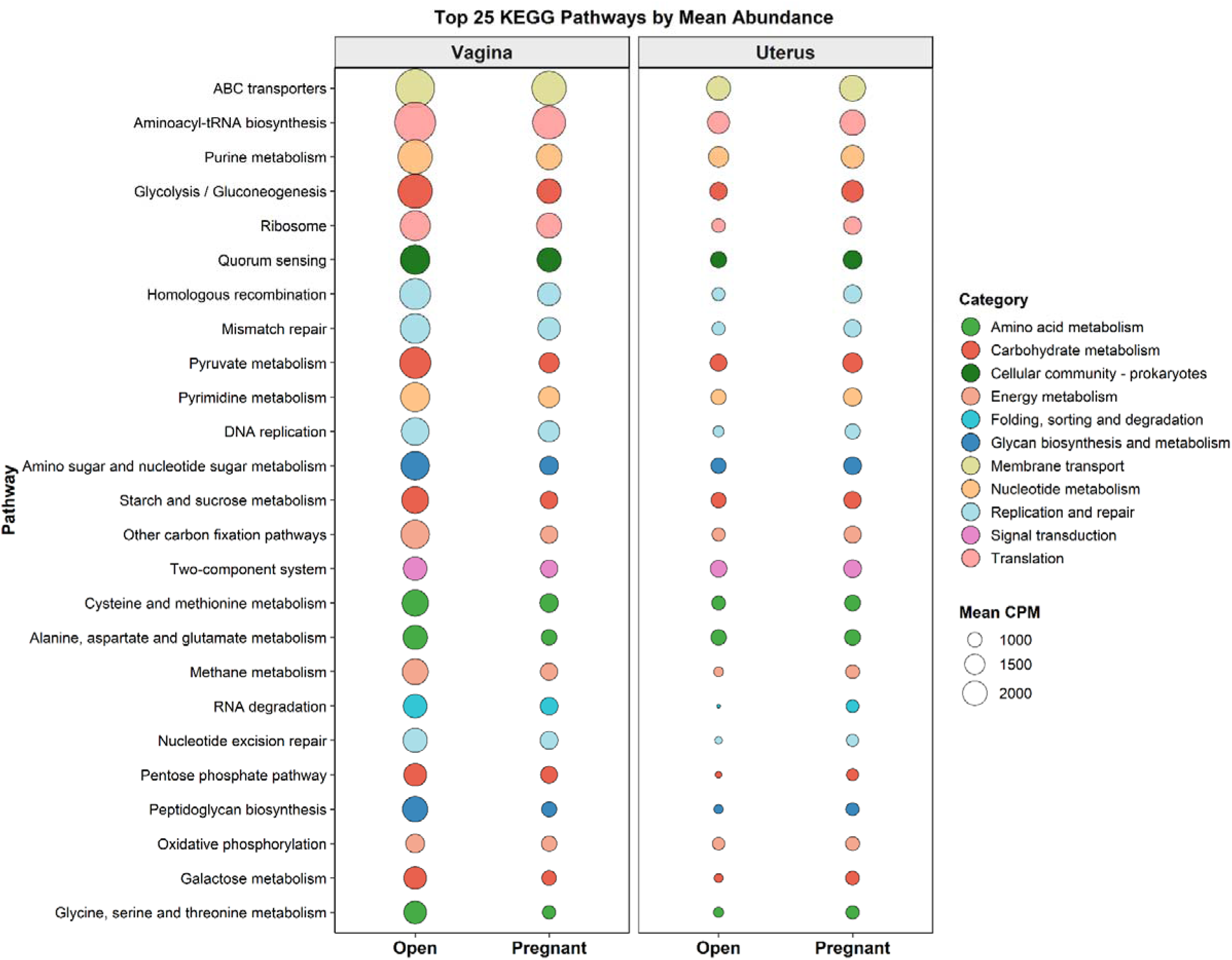
Functional potential of the vaginal and uterine microbiomes based on KEGG orthology (KO) analysis. Bubble plot showing KEGG pathways for vaginal samples (n = 53; pregnant = 47, open = 6) and uterine samples (n = 43; pregnant = 37, open = 6) by subsequent pregnancy status. Bubble size represents the mean pathway abundance expressed as counts per million reads (CPM), and color indicates the corresponding KEGG pathway category.

**Figure 5.**
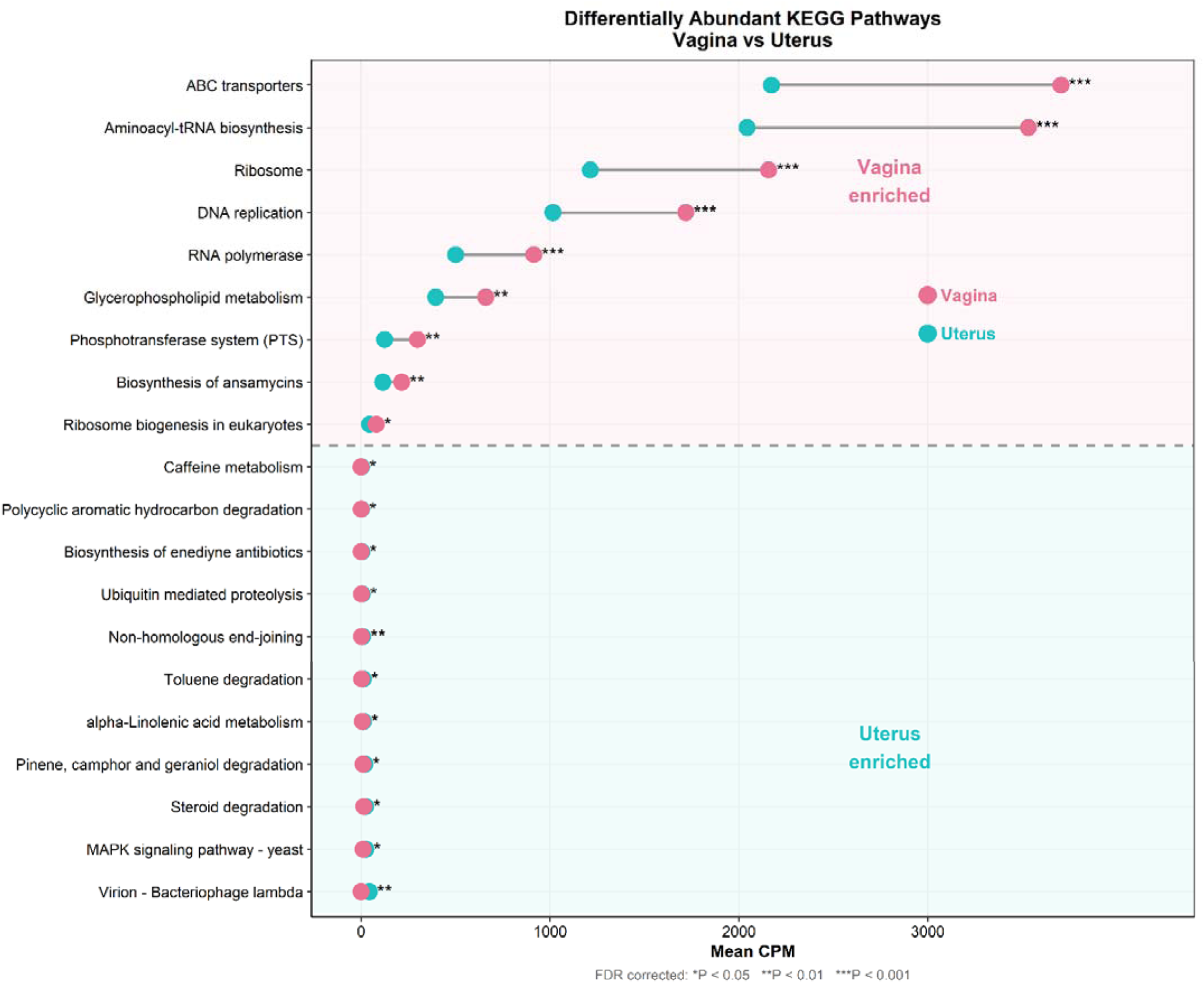
Lollipop plot illustrating the top 20 differentially abundant KEGG pathways between vaginal and uterine microbiomes of beef cows (vagina, n = 53 ; uterus, n = 43). Each dot represents the mean counts per million (CPM) for the vaginal (pink) or uterine (teal) samples, and connecting lines indicate the magnitude of difference in mean CPM between sample types. Pathways are grouped by site of enrichment, with vagina-enriched pathways shown in the upper pink-shaded section) and uterus-enriched pathways shown in the lower teal-shaded section, separated by a dashed line. Statistical significance was determined using the Mann-Whitney U test with Benjamini-Hochberg false discovery rate (FDR) correction. Significance levels: *P < 0.05, **P < 0.01, ***P < 0.001.

### Uterine microbiome

Similar to the vaginal samples, two uterine samples (D92 and D94) had no sequences assigned (≥ 10 reads) to a unique species. There were no differentially abundant genera or species in the uterine microbiome between open and pregnant cows (*P* > 0.05), and genus-level community structure also did not differ (Fig. 6A; PERMANOVA, *R²* = 0.009, *P* = 0.914). Likewise, genus-level richness and diversity indices at both genus and species levels were similar between groups (Fig. 6B-D). The uterine microbiome composition was dominated by the genera *Cutibacterium* (24.65%) and *Streptomyces* (20.45%), followed by *Staphylococcus* (4.17%), *Acinetobacter* (3.28%), *Alistipes* (3.27%), and *Methanocatella* (3.15%) (Fig. 7A). At the species level, the most abundant taxa were *Cutibacterium acnes* (31.63%) and *Streptomyces narbonensis* (26.19%), followed by *Caldifermentibacillus hisashii* (4.68%), *Brachyspira hampsonii* (2.98%), and *Staphylococcus chromogenes* (2.14%) (Fig. 7B). When stratified by reproductive outcome, dominant taxa were similar between open and pregnant groups, and no genera or species were differentially abundant between the two groups (*P* > 0.05).

**Figure 6.**
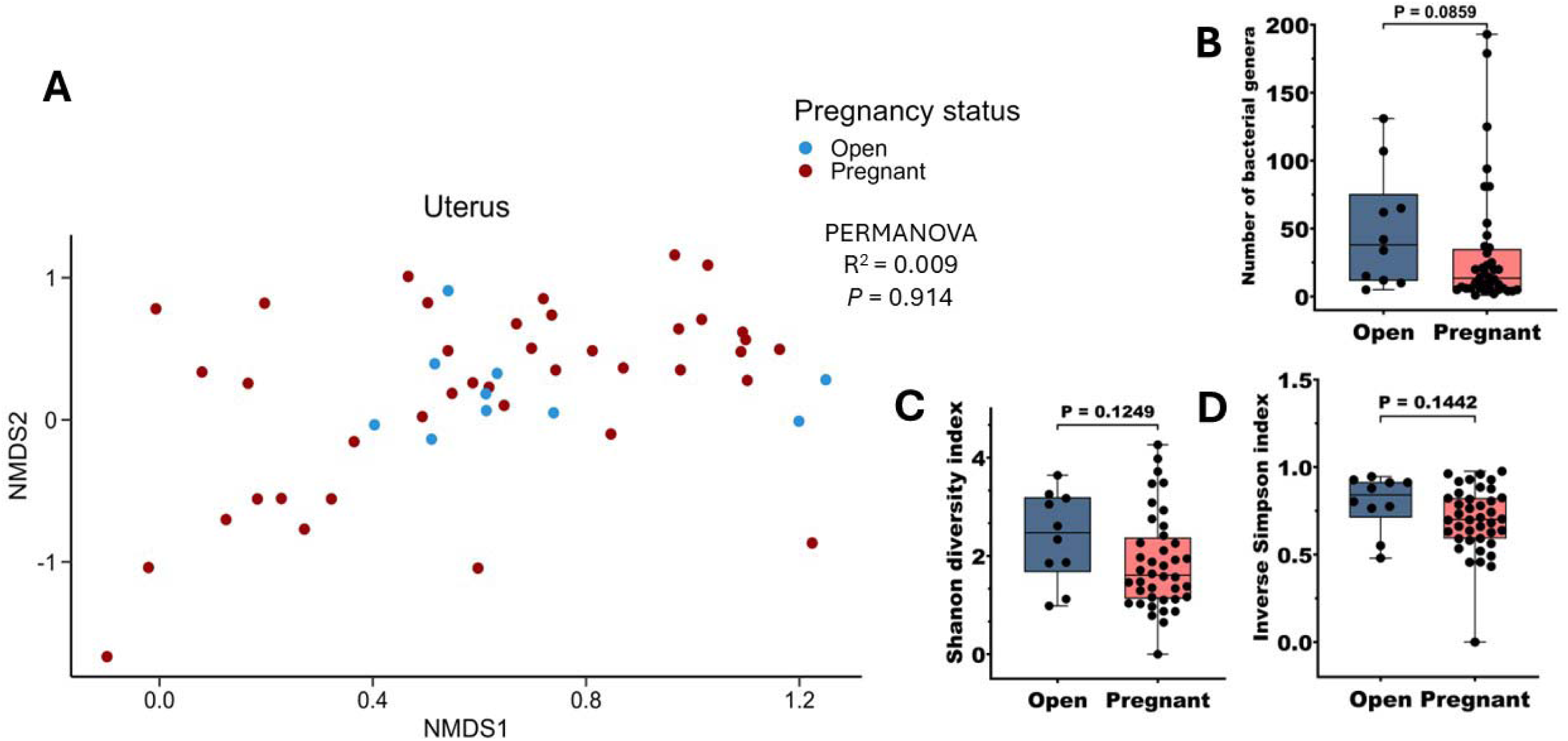
Alpha and beta diversity of the uterine microbiome in cows that remained open (n = 9) or became pregnant (n = 41) following artificial insemination (AI). Alpha and beta diversity of the vaginal microbiome at the genus level in cows that remained open (n = 7) or became pregnant (n = 53) following artificial insemination. (A) Non-metric multidimensional scaling (NMDS) plot based on Bray–Curtis dissimilarities calculated from genus-level relative abundances in vaginal samples from open and pregnant cows. Box-and-whisker plots showing alpha diversity metrics, including (B) observed genera richness, (C) Shannon diversity index, and (D) inverse Simpson diversity index. Boxes represent the interquartile range (IQR), the horizontal line indicates the median, and whiskers extend to 1.5 × IQR. Individual data points are shown as jittered dots. Statistical differences in genus richness and alpha diversity of the uterine microbiome between cows that remained open or became pregnant following artificial insemination were evaluated using the Mann-Whitney U test (two-tailed), with significance declared at P < 0.05.

**Figure 7.**
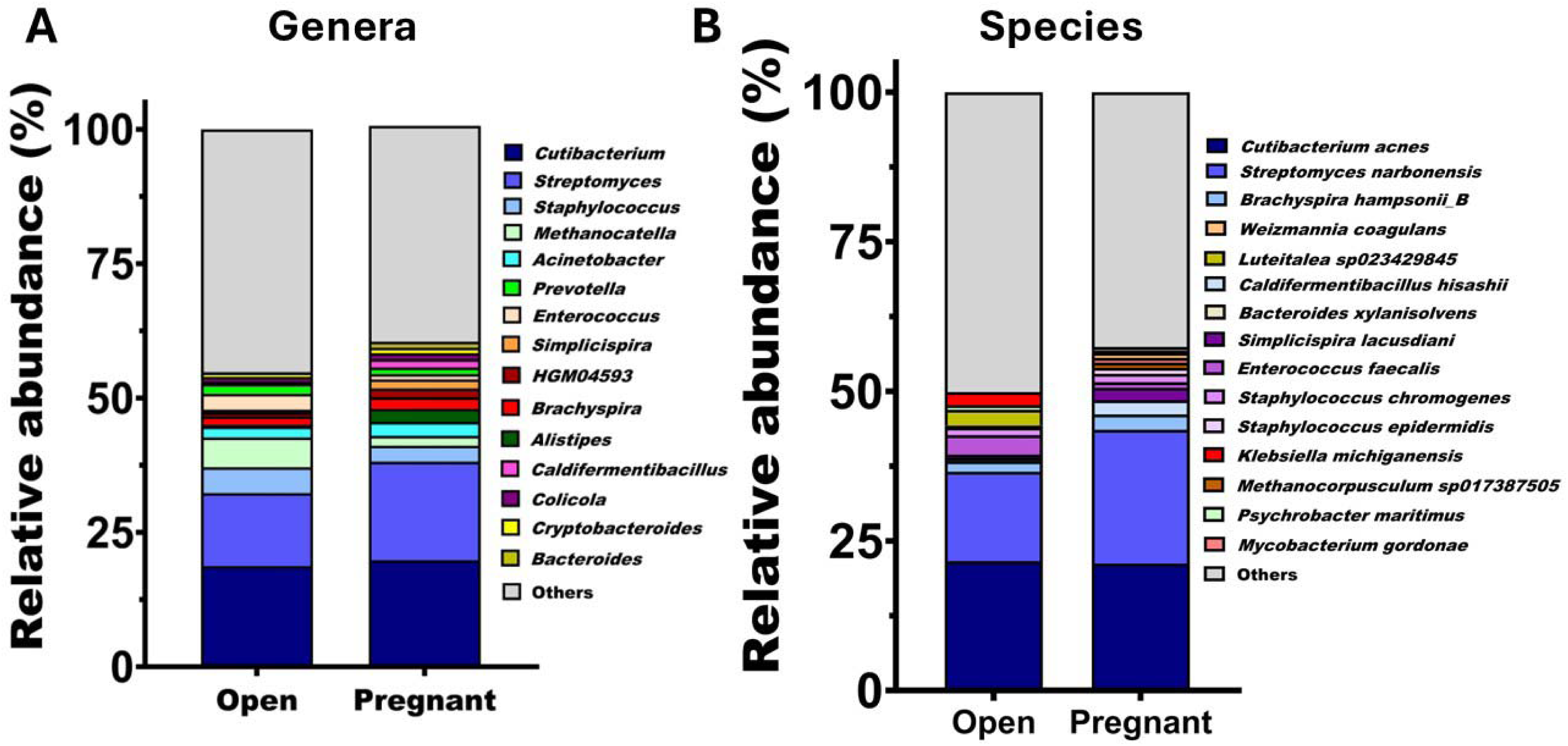
Stacked bar charts showing taxonomic composition of the uterine microbiome in cows that remained open (n = 9) or became pregnant (n = 41) following artificial insemination. (A) Relative abundance (%) of the 15 most abundant bacterial genera; (B) Relative abundance (%) of the 15 most abundant bacterial species.

The KEGG pathway analysis showed that no pathways differed significantly in the functional features of the uterine microbiome between open and pregnant cows (Fig. 4, Fig. S3B). The most abundant functional categories included nucleotide metabolism, membrane transport, replication and repair, carbohydrate metabolism, and energy metabolism. The dominant individual pathways included ABC transporters, aminoacyl-tRNA biosynthesis, purine metabolism, and glycolysis/gluconeogenesis (Fig. 4).

### Vaginal vs. uterine microbiome

Given that vaginal and uterine samples were collected simultaneously from the same animals, we compared these sites to assess niche-specific and shared microbial composition and functional potential across the reproductive tract. As expected, the vaginal and uterine microbiomes differed significantly at the genus level (PERMANOVA: *R²* = 0.20, *P* < 0.001; Fig. 8A). The uterus harbored greater taxonomic richness, with 471 genera and 707 species detected compared with 160 genera and 181 species in the vagina, of which 140 genera and 131 species were shared between the two sites (Fig. 8B, Fig. S2A-B). Alpha diversity indices were also higher in the uterus than in the vaginal microbiome (Fig. 8C-D). At the species level, these compositional differences were associated with a greater relative abundance of *S. pluranimalium* and *U. diversum* in the vaginal microbiome, and *S. narbonensis* in the uterine microbiome (MaAsLin3, FDR < 0.05).

**Figure 8.**
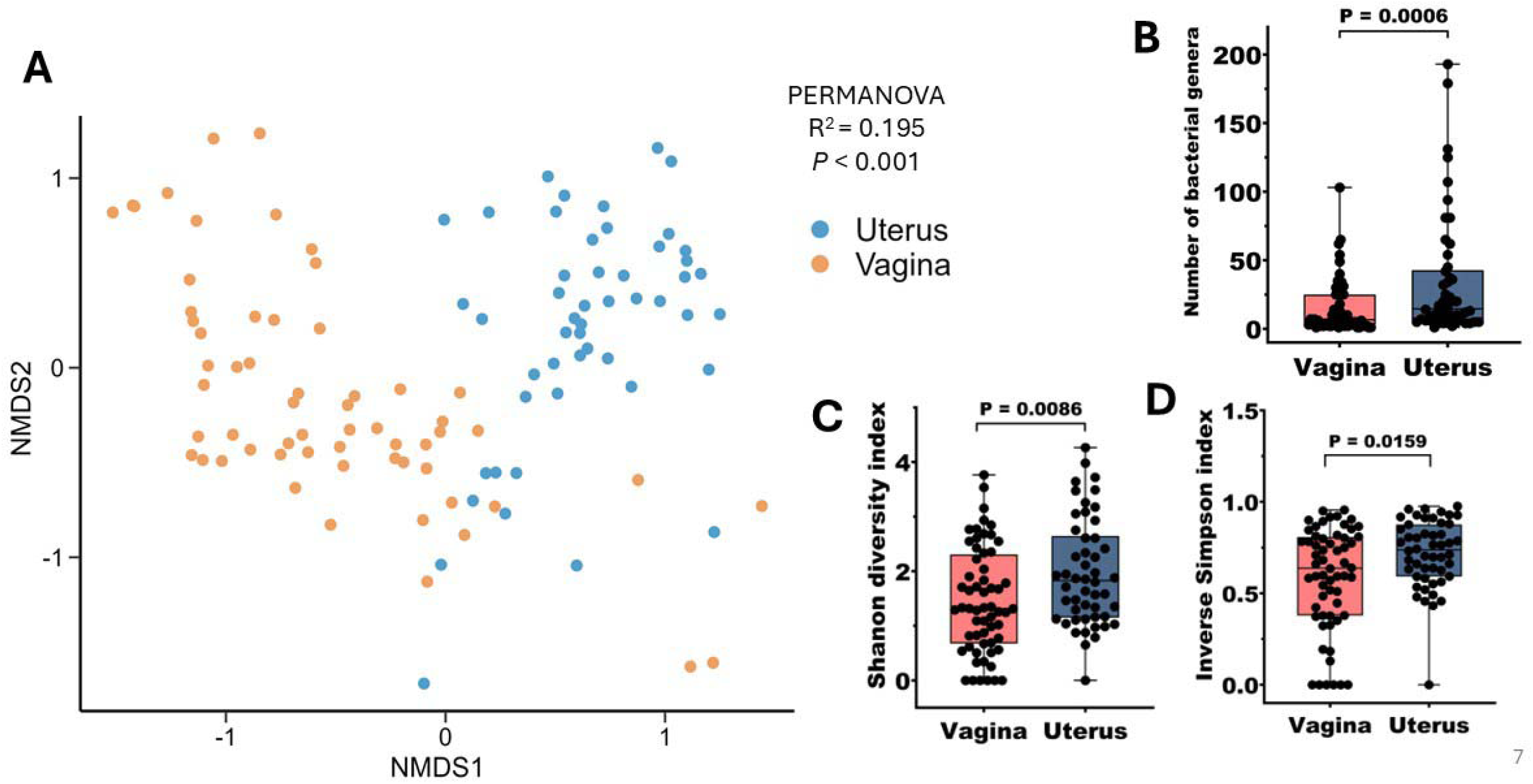
Beta and alpha diversity analysis of the vaginal. **(n = 60) and uterine (n = 50) microbiomes in cows that became pregnant or remained open following artificial insemination**. (A) Non-metric multidimensional scaling (NMDS) plot based on Bray–Curtis dissimilarities calculated from genus-level relative abundances in vaginal and uterine samples**(B),** Shannon index **(C),** inverse Simpson index **(D).** Boxes represent the interquartile range (IQR), the horizontal line indicates the median, and whiskers extend to 1.5 × IQR. Individual data points are shown as jittered dots. Statistical differences between the two anatomical sites were evaluated using the Mann-Whitney U test (two-tailed), with significance declared at p < 0.05.

Functional profiling of the uterine and vaginal microbiomes based on KOs showed no clear separation between cows that subsequently became pregnant and those that remained open following AI (Fig. S3A–B, Fig. 5). However, the functional composition of the uterine and vaginal microbiomes differed significantly (Fig. S3C; PERMANOVA: *R^2^* = 0.07; *P* < 0.001). There were 50 differentially abundant KEGG pathways between the vaginal and uterine microbiomes (FDR < 0.05), with 39 enriched in the vagina and 11 in the uterus (Fig. 5). Pathways enriched in the vaginal microbiome were primarily associated with genetic information processing, including ABC transporters, aminoacyl-tRNA biosynthesis, ribosome, homologous recombination, mismatch repair, and DNA replication (all *P* < 0.001), as well as quorum sensing and beta-lactam resistance. In contrast, the uterine microbiome showed enrichment in metabolic pathways including viral-associated and DNA repair pathways. At the broader functional category level, the vaginal microbiome exhibited higher relative abundance of pathways across most categories, with the largest differences observed for transcription (1.80-fold), translation (1.75-fold), and membrane transport (1.70-fold), whereas information processing in viruses was the only category more abundant in the uterus (Fig. 5).

### Antimicrobial resistance genes present in vaginal and uterine microbiome

A total of 105 ARGs spanning 11 antimicrobial classes were detected across 110 samples (vagina, n = 60; uterus, n = 50). The most abundant ARGs were primarily tetracycline resistance genes, namely *tet*(Q), *tet*(W), *tet*(O/W), *tet*(M), and *tet*(44). Other abundant ARGs included *lnu*(C) (lincosamides), *mel* (macrolides), and *bla*_TEM-116 (_beta-lactams) (Fig. 9A-B). ARG composition differed by anatomical site. The most abundant vaginal ARGs were *tet*(Q), *tet*(W), *lnu*(C), *tet*(M), and *tet*(O/W). In contrast, the uterine microbiome showed a more evenly distributed resistome, with ARGs conferring resistance to Macrolides, Lincosamides, and Streptogramin B (MLS_B_) (32.5%), tetracycline (21.3%), rifamycin (19.6%), and phenicol (7.2%) predominating. The most abundant ARGs in the uterine microbiome were *tet*(Q), *tet*(W), *bla*_TEM-116,_ *mel*, and *lnu*(C) (Fig. 9A, Fig. 10 A-B).

**Figure 10.**
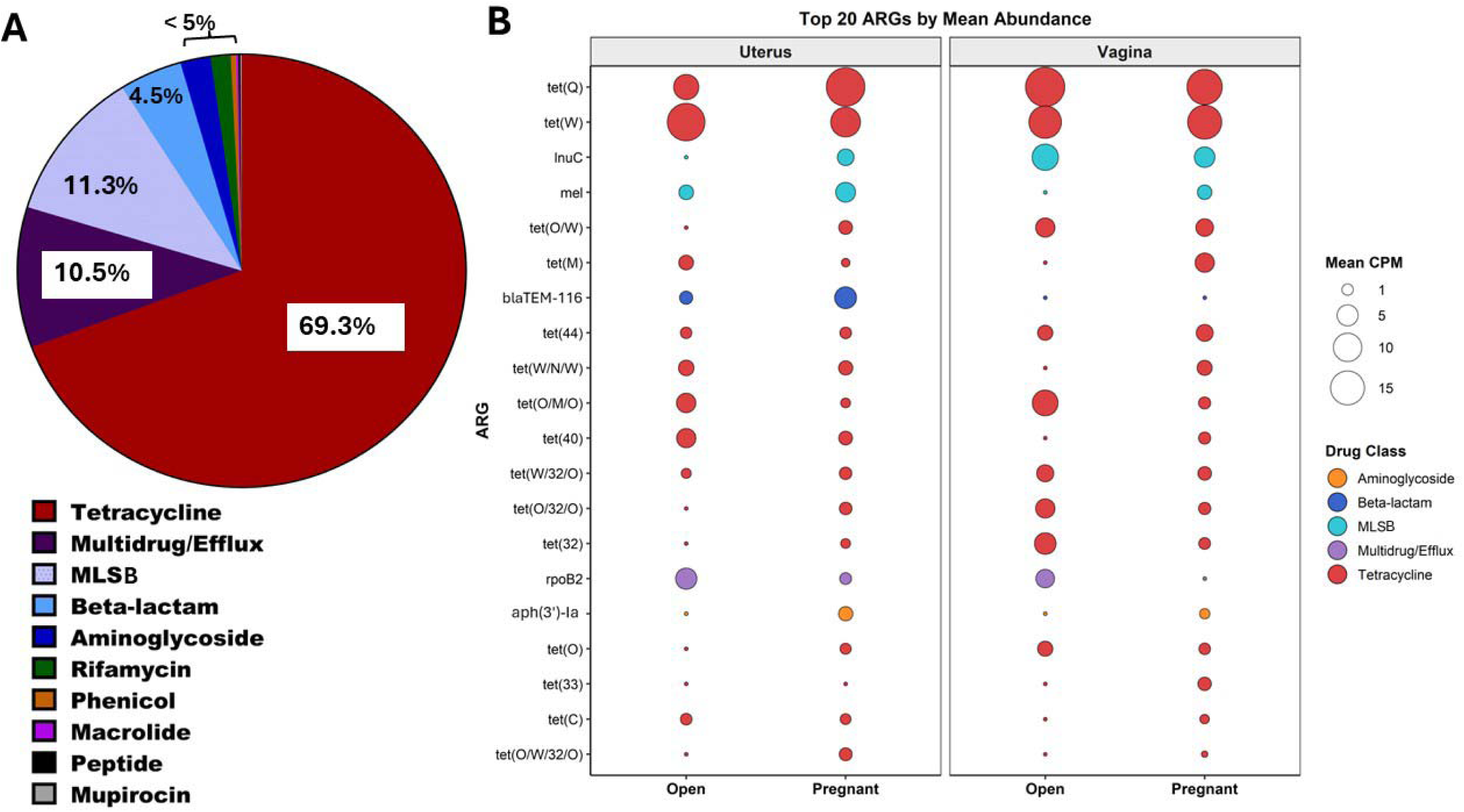
Antimicrobial resistance gene (ARG) profiles in bovine female reproductive tract microbiomes. **(A)** Pie chart showing the abundance (%) of ARG classes detected across all the vaginal and uterine microbiome samples. **(B)** Bubble plot of the 20 most abundant ARGs across vaginal and uterine samples from cows classified as open (non-pregnant) or pregnant following artificial insemination. Bubble size reflects mean ARG abundance in counts per million (CPM; scale: 1–15 CPM), and bubble color denotes the associated antibiotic drug class (legend, right). Dots indicate low-abundance detection below the CPM size threshold. ARGs are ordered by overall mean abundance (top to bottom).

**Figure 11.**
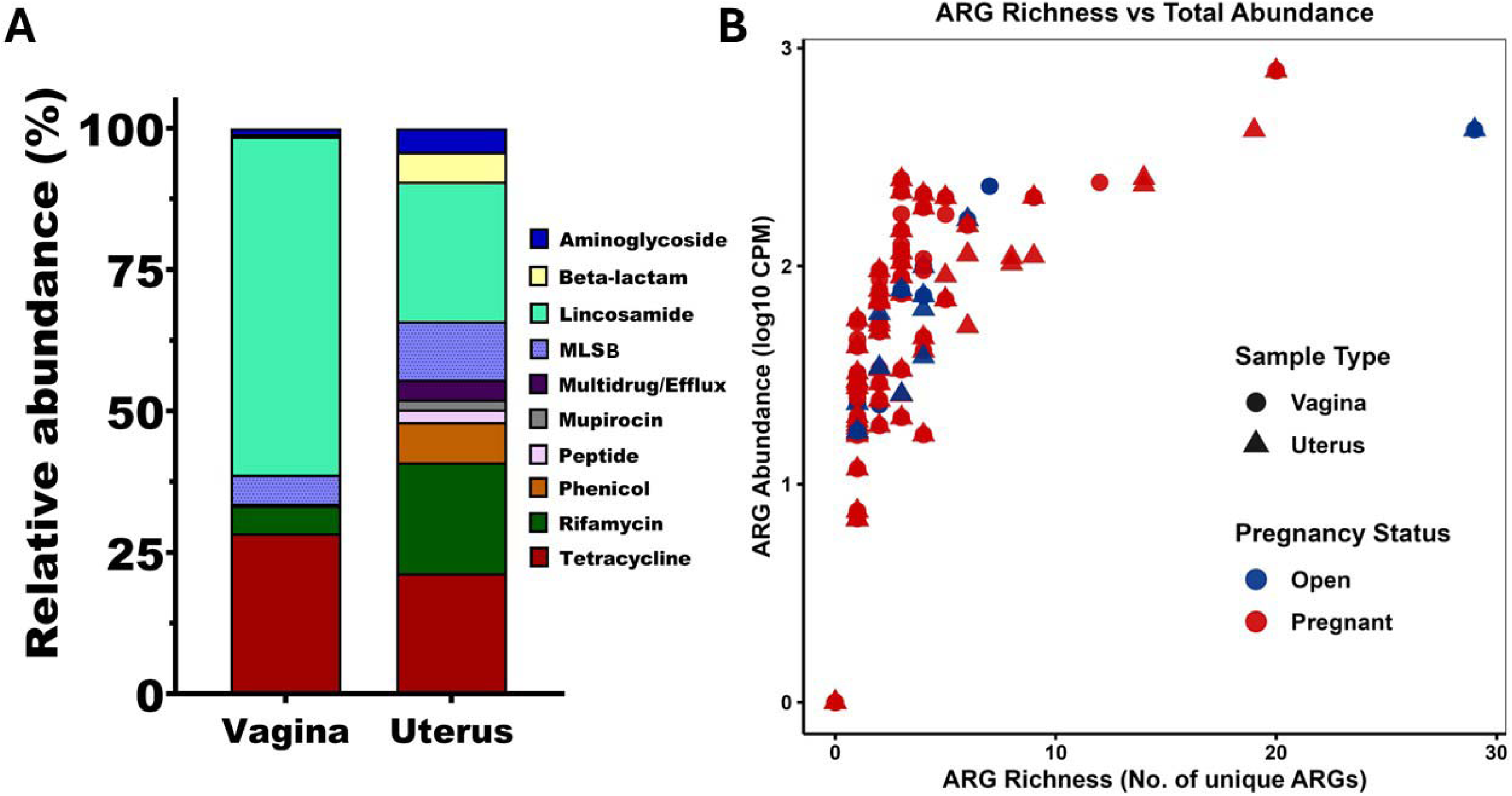
Composition and abundance of antimicrobial resistance genes (ARGs) in bovine reproductive tract microbiomes. **(A)** Stacked bar chart showing the relative abundance (%) of ARG classes detected in vaginal (n = 60) and uterine (n = 50) microbiomes from cows that became pregnant or remained open (non-pregnant) following artificial insemination. Stacked bars represent the proportional contribution of each ARG class across the two anatomical sites. **(B)** Scatter plot showing the relationship between ARG richness (number of unique ARGs per sample) and total ARG abundance (log_10_ counts per million reads, CPM). Each point represents an individual sample, with shape indicating anatomical site (circles = vagina; triangles = uterus) and color indicating pregnancy outcome (blue = open; red = pregnant). MLS: Macrolide–Lincosamide–Streptogramin resistance genes.

## DISCUSSION

There were no significant differences in vaginal microbiome structure associated with subsequent pregnancy outcome following AI. This result aligns with our previous 16S rRNA gene-based study, which also found no differences in the community structure of the vaginal microbiota in a similar cohort of mature cows (Webb et al., 2023). Similarly, Messman et al. (2020) reported that pre-breeding vaginal bacterial profiles did not significantly differ in diversity or community composition between cows that subsequently became pregnant and those that remained open following AI (Messman et al., 2020). Consistent with these findings, we also observed no differences in vaginal microbiome composition between outcome groups. However, cows that failed to conceive had significantly greater genus- and species-level richness and diversity in their vaginal microbiome compared to cows that subsequently became pregnant. This finding is consistent with our previous 16S rRNA gene sequencing study, in which richness (amplicon sequencing variants [ASVs]) and diversity tended to be greater (*P* = 0.05) in open cows compared to those that became pregnant (Webb et al., 2023). Thus, the current metagenomic study confirms a modest but significant increase in species-level diversity in the vaginal microbiome of open cows. This finding is also in line with observations by Laguardia-Nascimento (Laguardia-Nascimento et al., 2015), who reported that pregnant *B. indicus* heifers and cows displayed lower vaginal microbiome diversity than nonpregnant animals. One possible explanation for this observation is that a highly diverse or more evenly distributed vaginal microbial community may be associated with reduced likelihood of successful conception. For example, in humans, studies show that a less diverse vaginal microbiome, often dominated by *Lactobacillus* species, is associated with improved fertility outcomes, whereas higher diversity, as seen in bacterial vaginosis-like communities, correlates with reduced pregnancy rates (Koedooder et al., 2019b; Karaer et al., 2021; Baud et al., 2023a; Samama et al., 2025). Unlike humans, the bovine vaginal microbiome is characterized by a diverse community of facultative and obligate anaerobic bacteria rather than a *Lactobacillus*-dominated community (Adnane and Chapwanya, 2022, 2024). It is possible that a more even, species-rich community in open cows reflects a subtle dysbiosis or the absence of a “keystone” commensal that may support pregnancy establishment. Indeed, our previous work identified specific vaginal taxa (primarily Bacillota) that were enriched in heifers that failed to conceive, suggesting that particular taxa, or broader community structure, may influence the vaginal, uterine environment and sperm/embryo viability (Webb et al., 2023).

In the present study, the bovine vaginal microbiome at the time of AI was dominated by the genera *Streptococcus* and *Ureaplasma*, with the most abundant species being *S. pluranimalium* and *U. diversum.* These findings are consistent with previous 16S rRNA gene-based studies that identified Bacillota and Pseudomonadota, the phyla to which *Streptococcus* and *Ureaplasma*, respectively, belong, as core members of the bovine reproductive tract microbiome (Moore et al., 2017; Ault et al., 2019; Webb et al., 2023).

We did not detect any differentially abundant genera or species in the vagina between pregnant and open cows. This finding does not support our previous observations based on 16S rRNA gene amplicon sequencing, which identified several taxa differing by fertility status in heifers (Webb et al., 2023). This discrepancy may partly reflect methodological differences between sequencing approaches. The 16S rRNA gene sequencing may overrepresent certain taxa due to variable gene copy numbers across bacterial species (Kembel et al., 2012; Louca et al., 2018), whereas shotgun metagenomics captures a broader representation of microbial DNA including from taxa that PCR primers may amplify poorly.

The vaginal microbiome was functionally characterized by pathways spanning membrane transport, carbohydrate metabolism, translation, and microbial community signaling. ABC transporters ranked among the most abundant pathways, consistent with the nutrient acquisition and host cell interaction strategies documented for *U. diversum*, the second most abundant vaginal species identified in the present study (Santos Junior et al., 2021). Carbohydrate metabolism pathways, including glycolysis/gluconeogenesis and pyruvate metabolism, were also prominent; the *U. diversum* genome encodes enzymes of the Embden-Meyerhof-Parnas pathway, supporting its capacity for substrate-level energy production in the reproductive tract (Marques et al., 2016). Amino acid and purine metabolism pathways were also prominent, consistent with the metabolic requirements of *Streptococcus* spp., which rely on amino acid catabolism and de novo purine biosynthesis for growth and adaptation at mucosal surfaces (Willenborg and Goethe, 2016). As with the taxonomic analysis, no differentially abundant functional pathways were detected in the vaginal microbiome between pregnant and open cows.

Uterine microbiome composition at AI did not significantly differ between cows that became pregnant and those that remained open. Notably, shotgun metagenomic sequencing did not reproduce the findings of our earlier 16S rRNA gene amplicon sequencing study of cattle from the same farm, which identified 28 differentially abundant taxa in the uterine microbiota associated with pregnancy success (Webb et al., 2023). Similarly, (Ault et al., 2019) reported that uterine bacterial community composition immediately before AI differed significantly by pregnancy outcome, with samples from nonpregnant cows clustering more tightly relative to those from cows that subsequently became pregnant (Ault et al., 2019). The absence of detectable microbiome differences by fertility outcome does not necessarily refute a role for microbes in fertility; instead, it highlights the complexity and subtlety of host-microbe interactions in reproduction.

Shotgun metagenomic and 16S rRNA gene sequencing differ fundamentally in how they capture microbial diversity and abundance, and results are not always directly comparable (Quince et al., 2017; Durazzi et al., 2021; Lugli and Ventura, 2022). 16S rRNA gene-based studies are vulnerable to biases introduced by unequal library sizes, compositional constraints, and variable 16S rRNA gene copy numbers across bacterial species, all of which can complicate ecological and statistical interpretation (Kembel et al., 2012; Weiss et al., 2017; Louca et al., 2018). In contrast, metagenomic sequencing provides broader functional and taxonomic resolution but is particularly sensitive to challenges posed by host DNA contamination and low microbial biomass, which are especially pronounced in reproductive tract samples (Marotz et al., 2018; Rosen et al., 2020; Ahannach et al., 2021; Ong et al., 2021). Host-derived reads can comprise up to 95.9% of total reads in vaginal metagenomic datasets, and even after depletion, substantial host-derived contamination can persist, reducing the sensitivity of microbial detection (Ahannach et al., 2021). Together, these methodological differences likely contribute to the lack of agreement between approaches and highlight the importance of integrating multiple sequencing strategies and applying appropriate normalization methods in reproductive microbiome research.

Conception in cattle is multifactorial and largely influenced by the timing of ovulation, semen quality, and a receptive uterine environment (Chebel et al., 2004). Microbial community differences may exert modest effects or become more apparent at other time points, such as during implantation or early embryonic development (Gao et al., 2024). Because this study represents a single time point at breeding, transient or dynamic changes critical for pregnancy establishment may have been missed (Ault et al., 2019). Future longitudinal studies with larger, more balanced cohorts will be required to better resolve microbial features associated with pregnancy outcomes.

In the present study, we observed clear differences in microbial community structure between vaginal and uterine microbiomes, in agreement with previous studies (Moore et al., 2017; Ault et al., 2019; Webb et al., 2023). At the species level, the vaginal microbiome was dominated by *U. diversum* and *S. pluranimalium*, while the uterine microbiome was dominated by *C. acnes* and *S. narbonensis.* The diverse taxonomic composition observed across both anatomical sites likely reflects multiple microbial seeding routes characteristic of the bovine reproductive tract. Respiratory-associated taxa such as *Histophilus somni* and *S. pluranimalium* have been previously detected in reproductive tract samples of cattle and may reflect transfer via licking and grooming behaviors (Rodrigues et al., 2015; Messman et al., 2020; Macleod et al., 2026). Gut-associated organisms including *Bacteroides xylanisolvens, Enterococcus faecalis,* and *Brachyspira hampsonii* likely represent fecal inoculation, a recognized route of microbial transfer to the bovine reproductive tract given the anatomical proximity of the vulva to the anus (Sheldon et al., 2019). Skin-associated taxa such as *C. acnes, Staphylococcus epidermidis,* and *Staphylococcus chromogenes* may originate from contact with the animal’s own skin or that of herd personnel during routine handling and sampling (Taponen et al., 2008; De Visscher et al., 2014; Ahle et al., 2022). Environmental and aquatic organisms including *Psychrobacter maritimus*, *Paracoccus marinus*, and *Simplicispira lacusdiani* may reflect exposure to water sources (Zeineldin et al., 2017; Shi et al., 2019). Soil-dwelling taxa such as *S. narbonensis* are consistent with environmental exposure during grazing. The archaeal taxon *Methanocorpusculum sp.,* detected across both sites, likely reflects gastrointestinal origin, consistent with fecal inoculation as a route of microbial transfer to the bovine reproductive tract (Daquiado et al., 2014; Holman and Gzyl, 2019).

Collectively, these findings suggest that the bovine reproductive tract microbiome is shaped by a convergence of multiple ecological inputs rather than a single microbial source. Although strict sterile sampling techniques and contamination controls were employed throughout, residual background signal from reagent or environmental sources cannot be entirely excluded in low-biomass metagenomic datasets (Salter et al., 2014; Eisenhofer et al., 2019), and species-level interpretations should be made with this caveat in mind. These compositional differences likely reflect site-specific factors such as proximity of the vagina to the external environment, variations in pH and oxygen tension, and differences in local immune regulation (Sheldon et al., 2019). These findings are in agreement with Moore et al. (2017), who demonstrated that the bovine uterus (even in virgin heifers) contains a microbiome distinct from that of the vagina (Moore et al., 2017).

Microbial diversity was higher in the uterus than the vagina, a finding that contrasts with previous reports, given that the bovine uterus is typically considered a low-biomass environment. The cervix acts as a physical barrier that limits microbial ascent, thereby reducing bacterial diversity in the upper reproductive tract (Clemmons et al., 2017; Ault et al., 2019; Moreno et al., 2021). Previous 16S rRNA gene sequencing studies in cattle have reported lower uterine richness and diversity compared to the vagina (Webb et al., 2023; Winders et al., 2023). Similarly, Ault et al. (2024) reported greater phylogenetic diversity in vaginal samples relative to uterine body and horn samples in heifers. The higher uterine alpha diversity observed in the present study may, at least in part, reflect methodological differences in DNA extraction methods between sample types. As noted in the limitations, the standard DNeasy Blood and Tissue Kit failed to yield sufficient DNA from uterine samples, necessitating the use of a CTAB/phenol:chloroform protocol. Differences in extraction efficiency between protocols may have differentially influenced the detection of low-abundance taxa, potentially inflating apparent diversity estimates in the uterine samples. Additionally, dietary and physiological factors have been shown to influence the bovine uterine and vaginal microbiota (Winders et al., 2023; Luecke et al., 2024), and such variation may have contributed to the diversity patterns observed. Despite these differences, both sites contained overlapping and site-specific taxa. This overlap supports the concept of an ascending microbial continuum; however, the presence of site-specific taxa indicates that the uterine microbiome is not merely a subset of the vaginal microbiome and may reflect microbial transfer during calving or breeding (Clemmons et al., 2017; Ault et al., 2019).

Functional profiling revealed distinct metabolic signatures between the two anatomical sites. Vaginal microbiomes were enriched in pathways related to genetic information processing, including ABC transporters, aminoacyl-tRNA biosynthesis, ribosome function, homologous recombination, and DNA replication, which may reflect the metabolic demands of a microbially and immunologically active mucosal microenvironment (Amabebe and Anumba, 2018; Sheldon et al., 2019; Navarro et al., 2023). These functional annotations likely reflect broader microbial functional potential rather than the metabolism of specific substrates within the reproductive tract (Lee et al., 2015). Because functional profiles did not differ between pregnant and open cows, broad functional shifts detectable by KEGG pathway categories were not associated with fertility outcomes at the time of AI in the present study. However, the relatively shallow sequencing depth post-processing and unequal sample sizes between pregnant and open cows may have limited the detection of subtle differences in functional pathway abundance (Hillmann et al., 2018). Future studies integrating longitudinal sampling and host–microbiome interaction analyses will be necessary to determine how site-specific microbial dynamics influence fertility outcomes in cattle.

We detected 105 unique ARGs spanning 11 antimicrobial classes including tetracyclines, MLS_B_, beta-lactams, and aminoglycosides. Genes conferring resistance to tetracyclines [e.g., *tet*(W), *tet*(Q), *tet*(O), *tet*(M)] were the most abundant, which may reflect the widespread use of tetracyclines in cattle production (Xu et al., 2022; Messele et al., 2023). Across all samples, ARG richness was strongly and positively correlated with total ARG abundance (Spearman’s ρ = 0.888, *P* < 0.001), suggesting that samples with greater microbial biomass and sequencing depth captured a broader resistome. Several β-lactamase genes, including *bla*_TEM-116_, were enriched in uterine samples, likely reflecting associations with taxa present in the uterine microbiome (Hussain et al., 2021; Katonge and Ally, 2025). The detection of ARGs in clinically healthy cows suggests that the bovine reproductive tract may serve as a reservoir for ARGs, raising One Health concerns regarding potential dissemination to calves, bulls, or the wider environment (Fan et al., 2024). While culture-based studies have shown that most isolates remain phenotypically susceptible (Webb et al., 2023), these findings indicate that the reproductive tract may act as a reservoir for clinically relevant ARGs. As with other low-biomass metagenomic studies, sequencing depth post-processing likely limited the detection of low-abundance ARGs.

Metagenomic characterization of low-biomass mucosal surfaces in mammalian hosts presents inherent technical challenges(Hillmann et al., 2018; Ahannach et al., 2021; Gihawi et al., 2023), several of which were encountered in the present study and should be considered when interpreting the findings. Sequencing depth was a notable constraint across both sample types, as the low ratio of microbial to host DNA may have resulted in unclassified reads and potential under-detection of low-abundance taxa. Approximately 29% of bacterial reads from uterine samples could not be resolved to the species level, consistent with these challenges. Despite two rounds of host DNA removal and supplementation of the Kraken2 database with bovine and human reference genomes, residual host-derived reads may have persisted, a challenge widely acknowledged in metagenomic studies of mucosal tissues (Hillmann et al., 2018; Ahannach et al., 2021). Equimolar pooling of low-biomass libraries for metagenomic sequencing can introduce variability in read counts across samples, potentially contributing to the high proportion of unclassified reads observed in this study (Hillmann et al., 2018; Ahannach et al., 2021). Notably, different DNA extraction protocols were used for uterine and vaginal samples, and these differences in lysis efficiency and DNA yield may have contributed to variation and should be considered when interpreting comparisons between anatomical sites (Mattei et al., 2019). The imbalance between pregnant and open animals (7 vs. 53 vaginal; 9 vs. 41 uterine) may have reduced statistical power to detect subtle differences associated with successful conception (Weiss et al., 2017; Ault et al., 2019; Webb et al., 2023). Unequal group sizes can introduce variation in sequencing library size and further reduce statistical power (Weiss et al., 2017), a limitation also noted in previous bovine reproductive microbiome study (Webb et al., 2023). Future studies should employ deeper sequencing coverage, optimized host DNA depletion, and larger, more balanced cohorts with age and parity controlled as covariates to better characterize pregnancy-associated microbial signatures. Furthermore, the cross-sectional design restricted to a single time point at AI precludes assessment of temporal microbiome dynamics across the estrous cycle or during early gestation; longitudinal sampling designs will therefore be essential to address these limitations and more comprehensively characterize the vagino-uterine microbiome and its influence on bovine fertility.

## CONCLUSION

In summary, shotgun metagenomic sequencing revealed that the bovine vaginal and uterine microbiomes at the time of AI are compositionally and functionally distinct, yet no pregnancy-associated taxonomic or functional signatures were detected. However, cows that remained open exhibited greater vaginal microbial diversity compared to pregnant cows. Both urogenital sites differed in dominant taxa, functional pathways, and antimicrobial resistance gene profiles, with the uterus showing higher overall diversity. The detection of diverse antimicrobial resistance genes across both sites further indicates that the bovine reproductive tract may serve as a reservoir of resistance determinants. Future studies incorporating improved host DNA depletion strategies and larger, more balanced cohorts of pregnant and non-pregnant cows will be critical to enhance microbial resolution and identify subtle microbiome features associated with pregnancy outcomes in cattle.

## ACKNOWLEDGEMENTS

The authors would like to thank the managers and staff at the NDSU Animal Nutrition and Physiology Center and the Hettinger Research Extension Center for their invaluable help with animal husbandry and sampling. We also thank Kaycie Schmidt and Emily Webb for their assistance with sample collection.

## Author Contributions

S.A. and C.R.D. conceptualized and designed the study and oversaw its execution. Cattle management was performed by S.A., C.R.D., J.S.C., and K.K.S. Animal care and sample collection were carried out by S.A., C.R.D., and J.S.C. Sample processing was conducted by J.K., and S.A. Bioinformatics and statistical analyses were performed by D.B.H., S.A., and J.K. The manuscript was written by J.K. and S.A., and reviewed and edited by S.A., D.B.H., J.K., J.S.C., and C.R.D.

## DATA AVAILABILITY

All the sequencing data will be made available upon request from Dr. Samat Amat until they are deposited in NCBI at the time of publication of this manuscript.

## LIST OF FIGURES

**Supplementary Figure S1.**
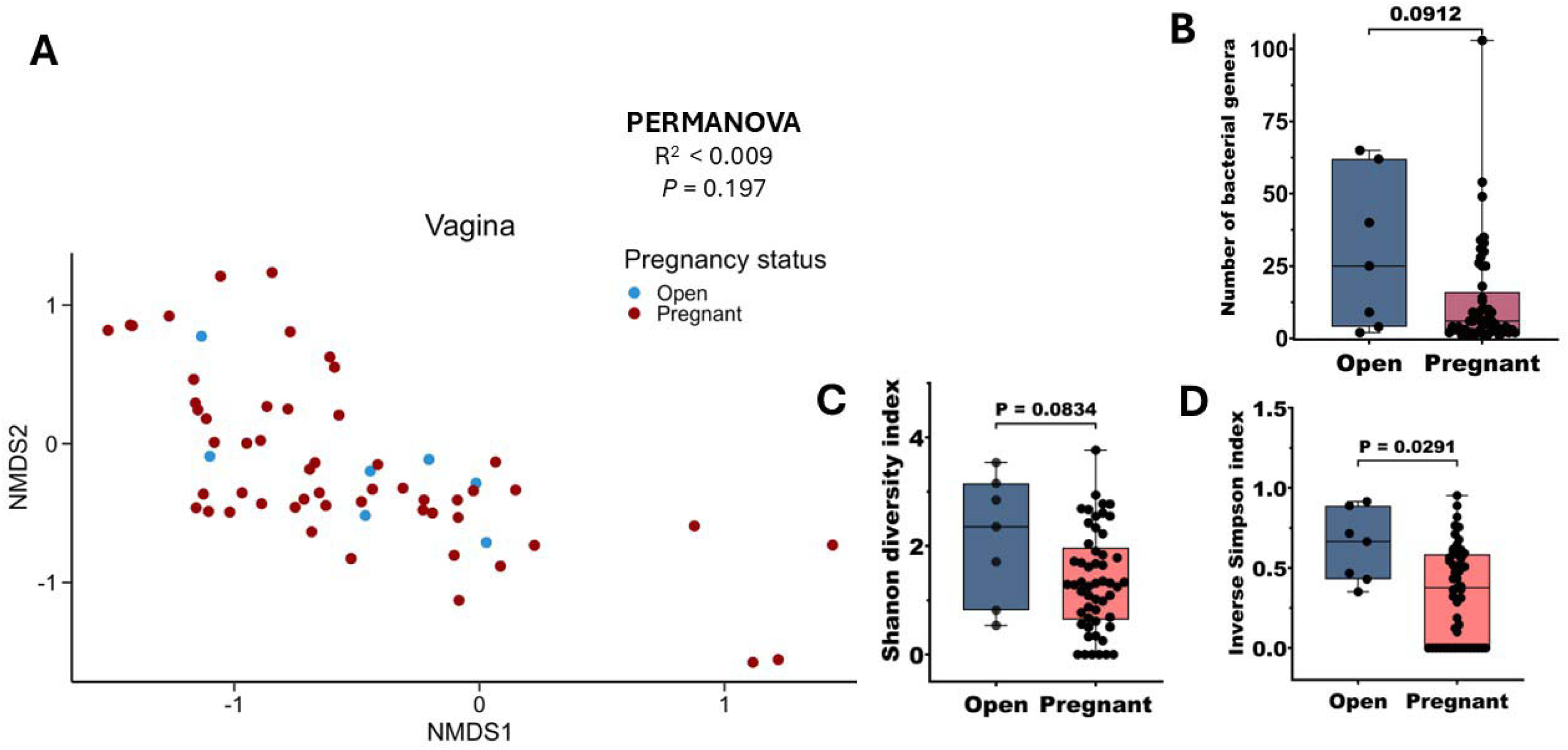
Alpha and beta diversity of the vaginal microbiome at the genus level in cows that remained open (n = 7) or became pregnant (n = 53) following artificial insemination. **(A)** Non-metric multidimensional scaling (NMDS) plot based on Bray–Curtis dissimilarities calculated from genus-level relative abundances in vaginal samples from open and pregnant cows. Box-and-whisker plots showing alpha diversity metrics, including **(B)** observed genera richness**, (C)** Shannon diversity index, and **(D)** inverse Simpson diversity index. Boxes represent the interquartile range (IQR), the horizontal line indicates the median, and whiskers extend to 1.5 × IQR. Individual data points are shown as jittered dots. Statistical differences in genus richness and alpha diversity of the vaginal microbiome between cows that remained open (n = 53) or became pregnant (n = 7) following artificial insemination were evaluated using the Mann-Whitney U test (two-tailed), with significance declared at P < 0.05.

**Supplementary Figure S2:**
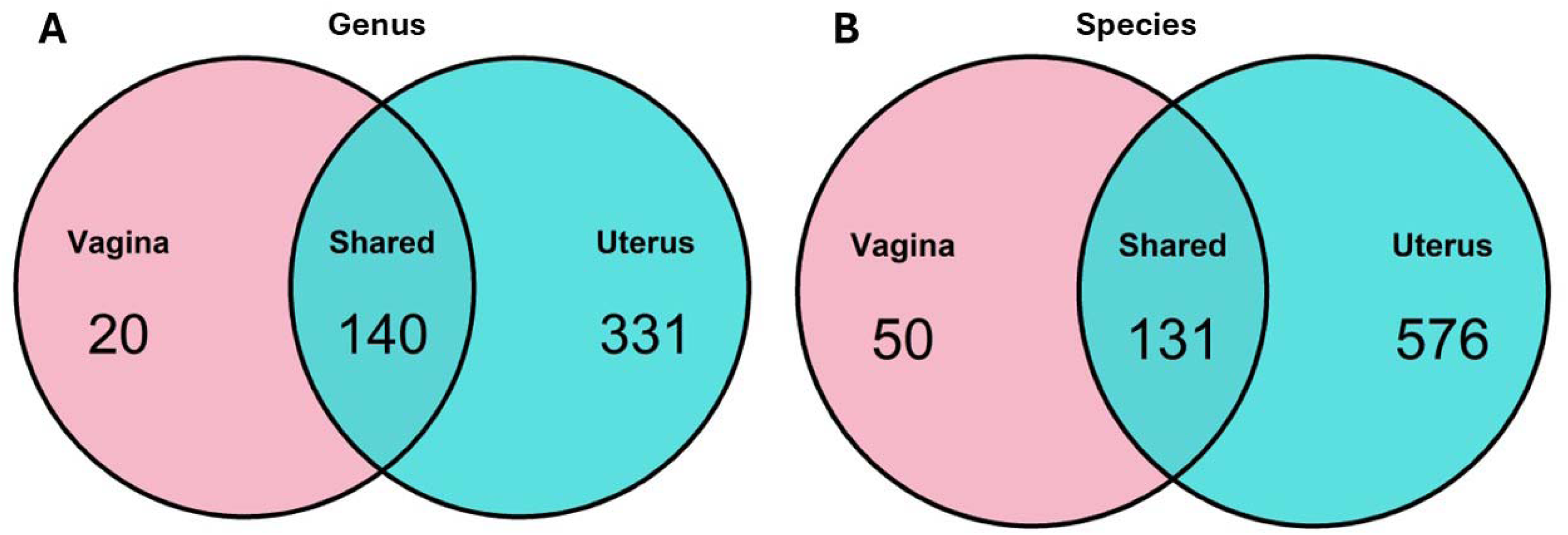
Overlap of bacterial taxa between vaginal and uterine microbiomes in cows following artificial insemination. Venn diagrams showing the number of shared and unique bacterial (A) genera and (B) species detected in the vaginal and uterine samples, irrespective of pregnancy outcome.

**Supplementary Figure S3.**
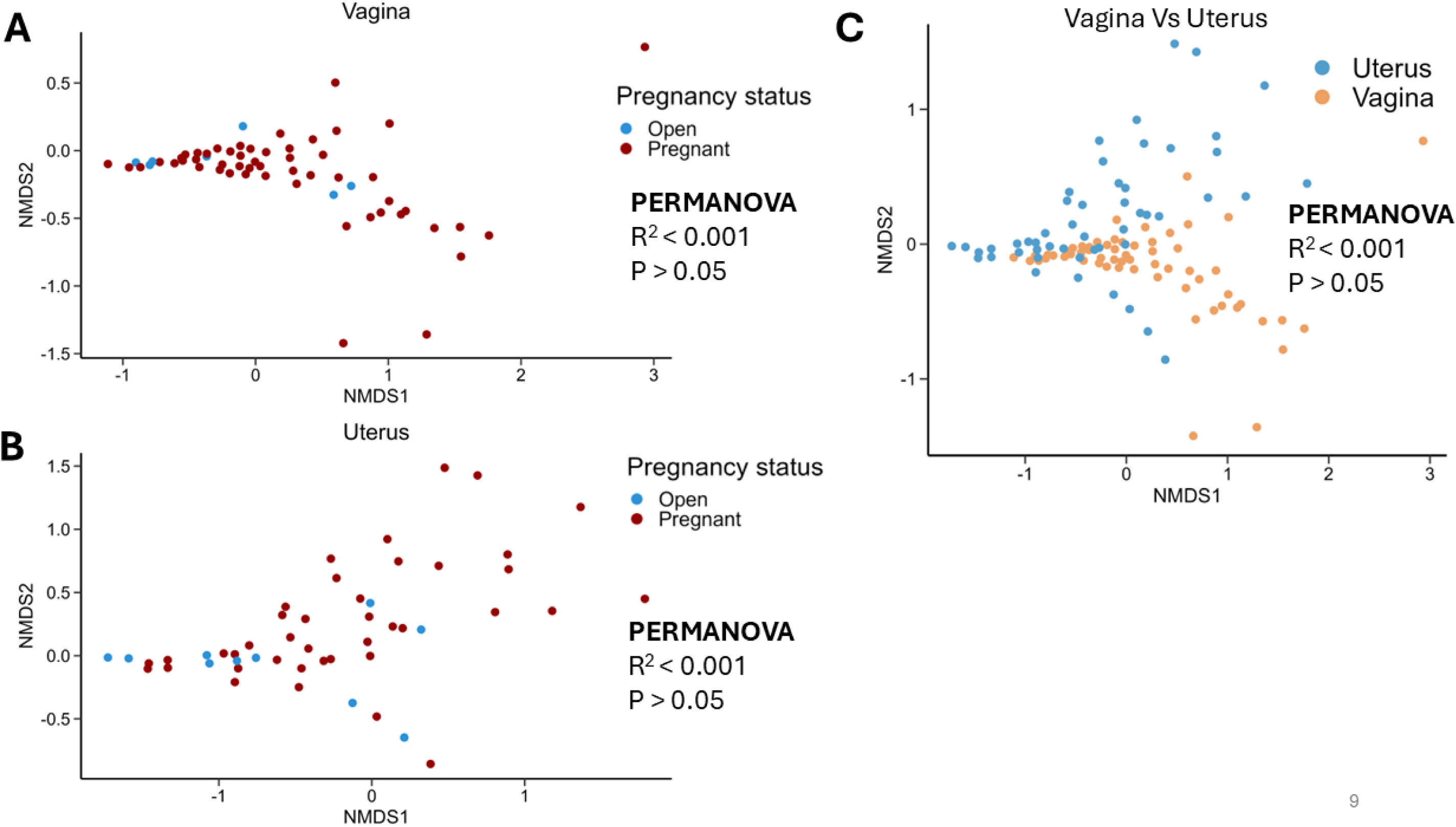
Non-metric multidimensional scaling (NMDS) ordination of functional profiles in bovine reproductive tract microbiomes. NMDS plots based on Bray–Curtis dissimilarities calculated from KEGG orthology (KO) counts per million reads (CPM) were used to evaluate functional differences between sample groups. **(A)** Vaginal samples (n = 53) from cows that became pregnant (n = 47) or remained open (n = 6) following artificial insemination. **(B)** Uterine samples (n = 43) from cows that became pregnant (n = 37) or remained open (n = 6). **(C)** Comparison of vaginal (n = 53) and uterine (n = 43) samples.

